# Prevalence of zoonotic hepatic nematode varies with small mammal community diversity across a heterogenous landscape in Eastern Uganda

**DOI:** 10.1101/2025.09.09.674808

**Authors:** Emilia Johnson, Diana Ajambo, Maria Capstick, Moses Arinaitwe, Olivia Ericsson, Fred Besigye, Jayna Raghwani, Tristan P W Dennis, Ronald Bogere, Andrina Nankasi Barungi, Aaron Atuhire, Candia Rowell, Namukuta Annet, Asmin Mohamed, Moses Adriko, Poppy H L Lamberton, Edridah Tukahebwa, Kathryn J Allan, Christina L Faust

**Affiliations:** University of Glasgow, School of Biodiversity, One Health, and Veterinary Medicine, Glasgow, UK; Ministry of Health, Vector Control Division, Kampala, Uganda; Royal Veterinary College, Pathobiology and Population Sciences, Hawkshead, UK; Liverpool School of Tropical Medicine, Department of Vector Biology, Liverpool, UK

**Keywords:** Calodium hepatica, zoonosis, land cover change, mitogenome

## Abstract

Identifying key drivers of pathogen infection prevalence and intensity in wildlife is essential to understand disease dispersal and transmission. *Calodium hepatica* (syn. *Capillaria hepatica*) is a generalist nematode that infects liver parenchyma of mammals worldwide and is capable of human infections. Prevalence ranges from 0-100% in wildlife, often varying across small geographic areas, making it an ideal parasite for understanding ecological drivers of variation. Here, we quantify prevalence of *Calodium hepatica* and present initial surveys of synanthropic small mammals in four villages representing differing land cover. Cross-sectional rodent trapping was conducted within and around households over consecutive dry seasons in Eastern Uganda. 18s rRNA gene of *C. hepatica* was amplified and a sub-set of PCR products sequenced to confirm presence of *C*. *hepatica.* Landscape structural diversity was classified by tree crown density and mean canopy height derived from 30m LiDAR data within a 0.5km buffer. Multivariable binomial generalised linear models were fit to *C. hepatica* prevalence. *C. hepatica* infection was common (overall 34.5%, CI95% 27.9-41.0) and found in rodent and shrew species inside and outside residences. We observe village-level differences in prevalence (18.2%–75.0%), with higher *C. hepatica* prevalence associated with higher relative proportion of *Rattus rattus* to other species (aOR=2.22 CI95% 1.30–3.85). Host diversity appears to be protective against parasite prevalence. Differences in molecular and macroscopy identification highlight challenges in diagnosis and a need for more specialized molecular tools. Further investigation is required to understand individual host and community variation in pathogen infection intensity and implications for zoonotic risk.

**Graphical Abstract:** 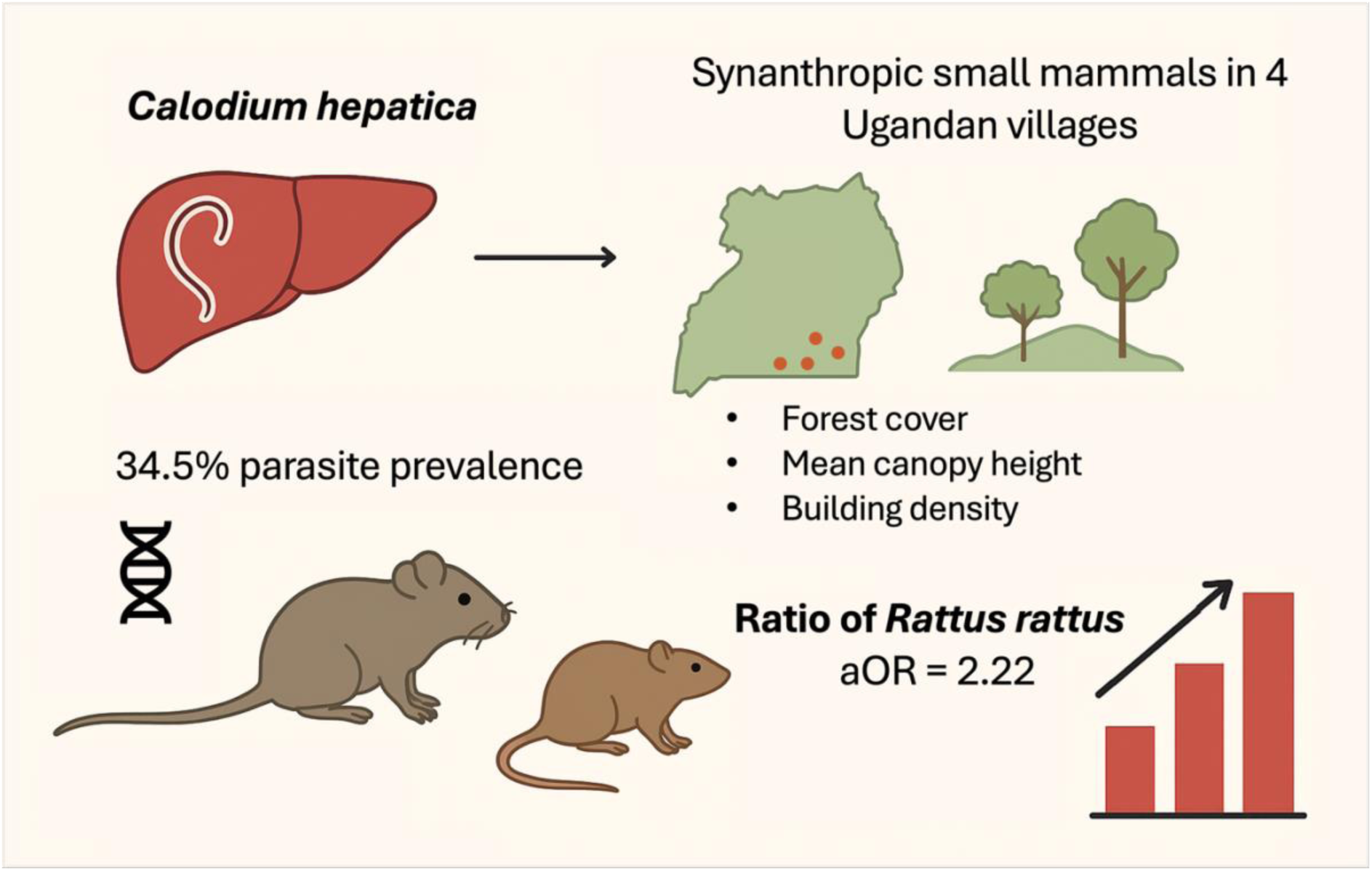

**Key Findings:**

- Zoonotic reservoirs, including *Rattus rattus*, *Mastomys erythroleucus* and *Crocidura olivieri,* are abundant in human-modified habitats and show elevated levels of household incursion
- High prevalence of *Calodium hepatica*, a nematode endoparasite, was identified in a range of small mammal species
- Higher proportion of native small mammal species relative to *Rattus rattus* appears protective against zoonotic pathogen load, with higher village-level *C. hepatica* prevalence in ecologically depleted sites
- Small-holder agriculture may provide a dilution effect through secondary wildlife support and small mammal competition, while dense village settings potentiate *C. hepatica* prevalence

## Introduction

Small mammals, including rodents and shrews, are reservoirs of zoonotic pathogens and play critical roles in the dispersal and transmission of infection. Rodents and shrews host pathogens related to over 60 diseases of public health concern, variably caused by bacteria, helminths, protozoans and viruses (Han et al. 2015, Mangombi *et al*., 2021). Land use change, for example urbanization or agricultural intensification, disrupts social and ecological systems and is regarded as a critical driver of pathogen emergence (Jones et al. 2013, LaDeau et al 2015). A key barrier to understanding emerging zoonotic disease is understanding the environmental conditions that link land use change, small mammal ecology and pathogen dynamics.

Many rodent-associated ectoparasites and endoparasites have implications for veterinary and public health (Isaac *et al*., 2018). Zoonotic rodent-borne diseases can be transmitted by ectoparasites, including fleas as the vector of plague (*Yersinia pestis*) (Adjemian *et al*., 2007) (Zimba *et al*., 2019) and tick-associated febrile diseases such as tick-borne relapsing fever (TBRF) and tularaemia (Dahmana *et al*., 2020). Rodent endoparasites can also contribute human or animal health problems, for example, contributing to the maintenance of *Toxoplasma gondii,* a parasite that causes disease in humans, cats, small ruminants and other mammals within an ecosystem (Dubey et al 2021). Small mammals have been shown to harbour diverse parasites in localised sites, for example in Nigeria, gross pathological investigation of endoparasites found small mammals to be infected with 12 helminth taxa (including *Strongyloides sp., Trichosomoides sp., and Trichuris sp.*) and 6 protozoan parasites (including *Plasmodium* sp. *Piroplasma* sp*., Toxoplasma gondii* and *Plasmodium sp.*) (Isaac *et al*., 2018) (Boundenga *et al*., 2019). Rodents are an important reservoir for human zoonotic schistosomiasis in West Africa (Catalano *et al*., 2018) and are an important refuge of *Schistosoma japonicum* in Southeast Asia, reducing public health gains made through treatment programs (Zou et al., 2020). Cataloguing and quantifying the burden of pathogens in wild rodent hosts is therefore crucial for informing interventions.

In addition to harboring important zoonoses, many small mammal species are highly adaptive, anthrophilic and thrive in human-altered landscapes. Global analysis has shown that the proportion of wildlife that host zoonotic pathogens increases with establishment of agriculture and land cover change, with particularly strong effect for rodent taxa (Gibb *et al*., 2020). Many rodent and shrew species have fast reproduction cycles and an early time to maturation, equating to rapid population turnover. Highly fluctuating populations and periods of high population density have been posited as a predictor of transmission risk and pathogen dispersal in rodent reservoirs, particularly for habitat generalist species (Ecke *et al*., 2022). In East Africa, invasive and native species of rodents and shrews occur in human-modified habitats in high abundances and host a diversity of zoonotic pathogens (i.e. Halliday et al 2013, Ogola et al. 2021, Rasoamalala et al. 2025). Understanding if specific host assemblages and associated land uses facilitate higher pathogen transmission is important for predicting disease risk and informing control strategies.

This study focuses on a generalist nematode parasite *Calodium hepatica* (syn *Capillaria hepatica*) in small mammal communities in Eastern Uganda*. C. hepatica* has an extremely broad host range: recorded in over 90 Muroidean rodent species and around 70 species of mammals, including carnivores, shrews, and primates (Fuehrer 2014 a,b). *Calodium hepatica* causes hepatic calodiasis (syn capillariasis) in heavily infected hosts. Adult worms develop in liver parenchyma and survive for approximately 6 weeks, with peak egg production at 40 days post infection (Fernandes et al 2001). Unembryonated eggs are released through fecal samples of infected hosts (Chieffi *et al*., 1891), but the hypothesized main route of release is through death or consumption of the infected host (Spratt and Singleton 2001). Eggs enter the environment through degradation of infected corpses or are mechanically passed through the gut of a predator (Momma 1930, Freeman and Wright 1960, Farhang-Azad 1977, Conlogue et al. 1979). To become infectious, the egg must embryonate in the environment, which is dependent on temperature and humidity (Luttermoser 1938, Pavlov 1955, Wright 1961, Spratt and Singleton 1986).. The embryonated eggs are then ingested by the rodent definitive host, or occasionally human hosts, then hatch and migrate to liver to continue the life cycle The parasite is primarily diagnosed through gross observations of lesions on the liver and morphological identification of eggs, but more recently PCR primers have been developed to identify infections (Williams et al 2018, Manor et al 2021).

*C. hepatica* has been found globally, but infection intensities and prevalence are highly variable. Typical of helminths, often there are a few heavily infected individuals with the majority having lighter infections (Conlogue *et al*., 1979, Reperant and Deplazes 2005, Sandy et al. 2024). Infection rates can be higher in mature or older hosts (Chieffi *et al*., 1891; Sinniah *et al*., 1979, Ceruti et al 2001, Rothenburger et al 2014) and has been associated with injuries, potentially associated with more aggressive individuals (Rothenburger et al 2014). Prevalence is positively correlated with small mammal host density in some studies (Davis, 1951; Childs *et al*., 1988; Meagher, 1999). Prevalence can range from 0-100% in populations on small spatial scales, even city blocks (Rothenburger et al 2014). Studies have also linked *C. hepatica* prevalence to habitat type (Davis, 1951; Childs *et al*., 1988; Roberts *et al*., 1992). Variation in prevalence across habitats has been hypothesized to be related to microclimate conditions affecting embryonation and transmission of eggs.

Human infection with *C. hepatica* has been recorded (Fuehrer et al 2011), presenting as a hepatic calodiasis associated with poor sanitation and low socioeconomic indices. Although only reported sporadically, lack of distinct clinical signs and appropriate diagnostics suggests human cases may be underreported (Marcelo et al 2010, Gonçalves et al. 2012). Inferring human risk from wildlife populations requires investigation of a myriad of complex features tailored to local ecological context. These include infection prevalence in reservoirs and host species abundance, key determinants of the force of infection that are prerequisites for transmission of pathogens into human populations (Lloyd-Smith *et al*., 2009) (Murray and Daszak, 2013). In vector-borne diseases, high prevalence of a pathogen in wildlife reservoirs has been linked to higher transmission risk to human populations (Johnson et al., 2023). For environmentally transmitted rodent-borne pathogens such as *Leptospira,* high household ‘rattiness’ – a composite proxy for rodent abundance – was found to be the factor most associated with higher infection risk for human residents (Eyre *et al*., 2022). However, further statistical modelling found that environmental variables such as vegetative land cover and household flooding were more influential than either rodent abundance or individual rodent pathogen shedding for human transmission risk (Soni *et al*., 2024). Other factors, such as infectious dose and human-wildlife contact patterns, will also be essential to understand transmission but fall outside the scope of this study.

Here we describe a survey of *C. hepatica* infection in small mammals conducted in eastern Uganda across a heterogenous of landscape of increasing human population and concomitant changes in land cover composition. Our study aimed to quantify infection prevalence and investigate the relationship with host factors (species, diversity, relative abundance) and landscape factors (forest cover, building density) to understand local drivers of infection.

## Materials and methods

### Site selection and household-level data

This study focussed on infection of *Calodium hepatica* among small mammals in sites in Eastern Uganda, within 33° 17’ 0.49255" N, 0° 13’ 34.6728"W and 33° 58’ 40.7611056006" N, 0° 44’ 4.9775928003" E. Climate is mainly tropical, characterised by bimodal wet seasons typically occurring from March-May and September-November (Kizza *et al*., 2009).

Live trapping was carried out in four villages across two dry seasons in July 2018 and February 2019 in Mayuge and Bugiri Districts, Eastern Uganda (Figure 1). Villages ranged in density of households and forest cover and loss (Table S1) and were chosen to represent a spectrum of land uses: fishing boat landing sites (Bugoto), small holder agriculture (Musubi), and timber plantations (Walumbe, Waka Waka). All sites had agricultural land and were adjacent to Lake Victoria but differed in the primary land cover and use. Trapping and sampling protocols were reviewed and approved by the Vector Control Division Research Ethics Committee (VCDREC095), Ugandan National Council for Science and Technology (NS-622) and the University of Glasgow’s School of Veterinary Medicine Ethics (27a/18). A minimum of 10 households were recruited from each village. In 2018, questionnaires were administered to households to ascertain observed levels of rodent infestations (Table S2).

**Figure 1.**
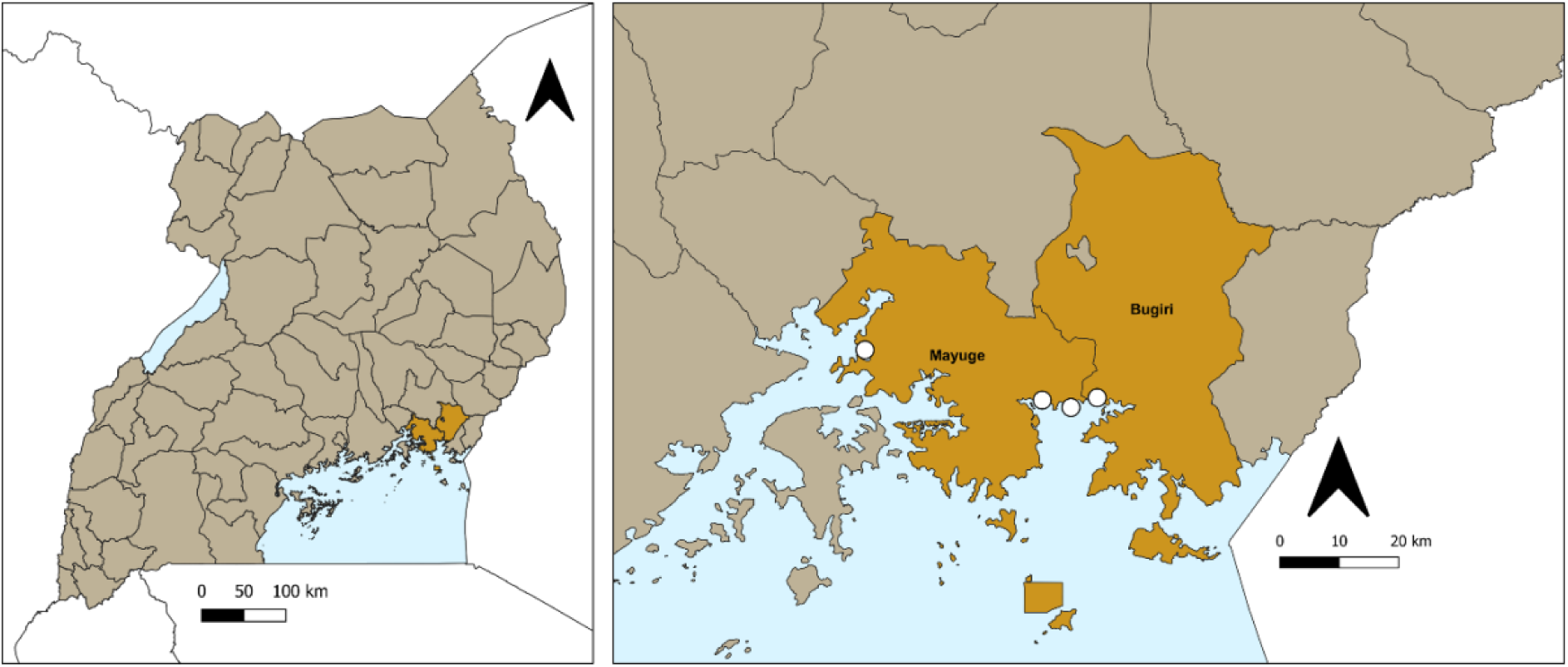
Map of study locations. (Left) Mayuge and Bugiri Districts in Eastern Uganda. **(Right)** Study sites (white points) in Mayuge and Bugiri, adjacent to Lake Victoria.

Within each household, a target of three traps were placed indoors and two traps were placed outdoors. A mixture of large Sherman traps (HB Sherman Traps, Tallahassee, USA. Dimensions: 7.6 x 8.9 x 22.9 cm; n=80), medium Tomahawk (Tomahawk Live Trap, Hazelhurst, USA. Model 602; dimensions 12.7 x 12.7 x 40.6 cm; n=20) and local handmade treadle traps (here called Tomahawk-sized treadle traps; multiple captures; approximate dimensions 15 x 13 x 40 cm; n=6) were used. Placement and number of traps was modified based on homeowner agreements. For each household, traps were left open and baited for three to five consecutive nights and checked first thing in the morning. Full traps were taken to a central processing unit for sample collection. Actual sampling effort varied between households, villages, and sessions due to logistical constraints.

### Individual specimen collection

Full traps were taken to a central processing area for humane euthanasia and postmortems. Individuals were euthanized via chloroform overdose and confirmation of death was by cervical dislocation. Morphometric measurements (head-body length, tail length, right ear, right hind foot) and weight were taken prior to dissection. All liver lobes were inspected for lesions and presence of *Calodium hepatica*. Liver tissue was also stored in 70% ethanol with nuclease-free water for downstream laboratory analysis.

### Environmental classification

Satellite-derived remote sensing datasets were used to assemble local landscape features and human metrics (Table 1). Gridded UN-adjusted human population estimates were assembled at 1 km resolution (WorldPop, 2018). Counts of building footprints were extracted from Google Open Buildings at each site as a proxy for household density and urban environments (Sirko *et al*., 2021). Elevation data was obtained from NASA SRTM 90 m Digital Elevation Database v4.1 (CGIAR-CSI) (Jarvis, A., H.I. Reuter, A. Nelson, 2008) and broad climatic trends (min/max temperature, precipitation) were assembled from BioClim (Fick and Hijmans, 2017). Given extensive conversion of agricultural land, tree cover was derived from Hanson’s Global Forest Watch (30 m) (Hansen *et al*., 2013) contemporaneously and with a time lag of 2, 5 and 10 years prior to trapping. Classification of tree cover was examined at multiple thresholds, ranging from conservative (≥50% of crown density per pixel) to modest thresholds (≥30). Mean canopy height (metres) was derived from Global Ecosystem Dynamics Investigation (Potapov *et al*., 2021a) using LiDAR measurements at 30m resolution. Proportion of cropland in 2019 was extracted from a dataset of global cropland expansion derived from Landsat time-series data at 30m resolution (Potapov et al 201b). To protect individual household identity, village centroids were taken as the mean coordinates of geolocated households and used to estimate spatial data. Environmental and anthropogenic covariates were extracted around village centroids within buffer radii of 0.5, 1, 2, 5, and 10km.

**Table 1.**
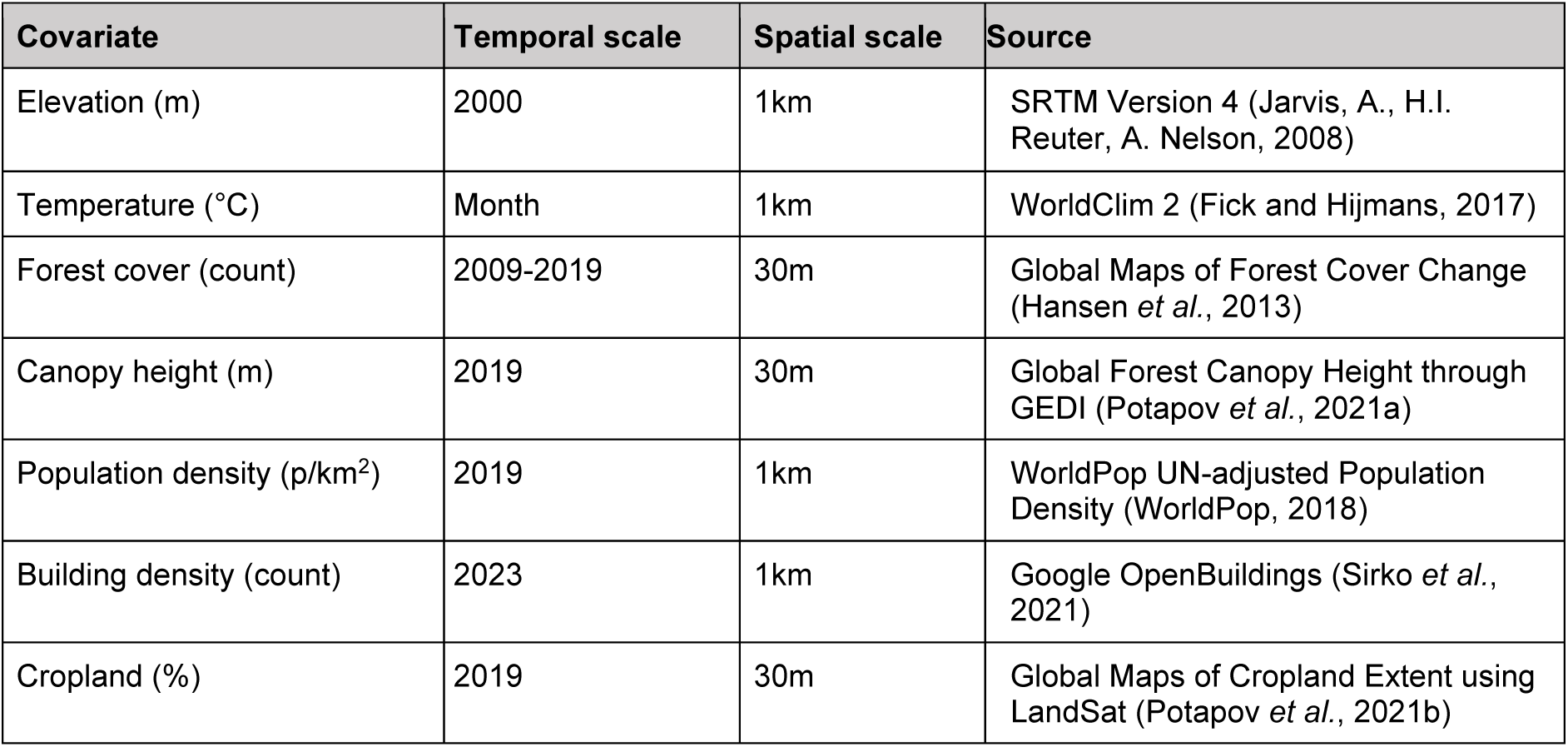
Summary of environmental and anthropogenic covariates scales and source.

| Covariate | Temporal scale | Spatial scale | Source |
| --- | --- | --- | --- |
| Elevation (m) | 2000 | 1km | SRTM Version 4 (Jarvis, A., H.I. Reuter, A. Nelson, 2008) |
| Temperature (°C) | Month | 1km | WorldClim 2 (Fick and Hijmans, 2017) |
| Forest cover (count) | 2009-2019 | 30m | Global Maps of Forest Cover Change (Hansen *et al.*, 2013) |
| Canopy height (m) | 2019 | 30m | Global Forest Canopy Height through GEDI (Potapov *et al.*, 2021a) |
| Population density (p/km<sup>2</sup>) | 2019 | 1km | WorldPop UN-adjusted Population Density (WorldPop, 2018) |
| Building density (count) | 2023 | 1km | Google OpenBuildings (Sirko *et al.*, 2021) |
| Cropland (%) | 2019 | 30m | Global Maps of Cropland Extent using LandSat (Potapov *et al.*, 2021b) |

### Laboratory analysis

DNA was extracted for host species confirmation and *C. hepatica* molecular diagnosis. DNA was extracted from 25mg of liver tissue with QIAgen Tissue and Blood Kit (Qiagen Ltd. Hilden, Germany) using manufacturer’s instructions with a 90-minute incubation with proteinase K and eluted into a final volume of 200μl. Extracted DNA was quantified using Qubit Broad Range Assay kits (Thermo Fisher Scientific, USA). For species, host cytochrome b (cytb) DNA was amplified using established protocols (Schlegel *et al*., 2012). Bands were visualised on 1.0% agarose gels and bands matching the ∼900 bp band were shipped to the UK and submitted for Sanger Sequencing with forward primers. A subset were also submitted with reverse primers. Resulting sequence traces were checked for clarity and trimmed with UGENE (Okonechnikov *et al*., 2012).

To identify presence/absence of *C. hepatica* DNA, the 18s rRNA gene of *C. hepatica* gene was amplified using previously published primers (Williams *et al*., 2020). PCR conditions were adapted and optimised for MgCl2, primer concentration, and thermal profile to increase specificity and sensitivity. The final PCR thermal profile was an initial denaturation at 94°C for 5 minutes, then 35 cycles (denaturation at 94°C for 30 seconds, annealing at 59°C for 30 seconds, extension 72°C for 30 seconds), then final extension at 72°C for 2 minutes. Final PCR was performed using the following master mix in 20μl total reaction volumes: 10x PCR buffer, 1.5mM MgCl2, 1U Taq DNA polymerase (Invitrogen™ 18038026), 0.2mM of each dNTP (NEB™ N0447S), 200nM forward primer, 200nM reverse primer, template DNA.

Initial PCR screening with liver DNA revealed a high incidence of very faint PCR bands (n =29/203). Several of these samples had observable parasites during necropsy (Dataset 1). As liver tissue can have a high proportion of inhibitors that can affect amplification of DNA (Al-Soud et al 2005), DNA was diluted 1:10 in nuclease-free water and PCRs were re-run. A sub-set of PCR product from strong bands (n =8) were sent for sequencing at DNA Sequencing and Services, MRC PPU, to confirm presence of C. *hepatica.* As both PCR reactions resulted in very faint bands, we conservatively estimated molecular presence of *C. hepatica* as a strong signal on either undiluted or 1:10 diluted DNA or a faint signal on both. Individuals with a faint signal on only one protocol were considered to be false positive. This likely underestimates prevalence but was decided to be the most specific diagnostic.

To characterize genetics of *C. hepatica* present in the region, DNA was extracted from a whole *C. hepatica* worm dissected from a *Rattus rattus* from Waka. DNA extraction followed manufacturer’s instructions for Beckman Coulter 280 Agencourt DNAdvance gDNA purification kit and single cell Illustra GenomiPhi DNA amplification kit (GE Healthcare Life Sciences, USA). DNA was submitted for sequencing through Novogene on an Illumina NovaSeq 6000, aiming for 2M paired-end reads, as no reference genome exists this was targeted to attempt to reassemble the mitochondrial genome.

### Data analysis

Adjusted trap success was used as a measure of relative abundance of small mammals for each village in each trapping session, calculated as total animals caught (n) as a proportion of trap nights corrected for misfired traps (number of traps set x number of nights – sprung traps * 0.5) (Nelson and Clark, 1973) and compared between village, household and trapping method. To assess whether sampling effort was sufficient to characterise host diversity, species accumulation curves were calculated for each site, for each trapping season, and overall. In addition, the proportion of households with small mammals was calculated as a general measure of household infestation. Household trap success was compared to qualitative surveys on household rodent control practices and lived experience of rodent contact. Two-sample T-tests were used to compare the adjusted trap success and proportion of households with small rodents between the two study districts.

Prevalence of zoonotic hepatic nematode *C. hepatica* were then calculated per village per year as the overall number of rodents or small mammals with molecular confirmation of *C. hepatica* (according to above diagnostic criteria) out of number tested by molecular methods. Binomial confidence intervals for point prevalence estimates were calculated using the package DescTools and used to compare the proportion of *C. hepatica* infection between species, and between sex within-species. Statistical analysis was performed in R (R Core Team, 2021)

To assess village-level difference in prevalence, binomial confidence intervals at a 95% threshold were calculated around point estimates. To assess possible environmental or human correlates of prevalence, bivariable analyses were conducted independently for each explanatory variable within a 0.5km buffer. Coefficients from binomial generalised linear model (GLM) were fit and coefficients exponentiated to calculate odds ratios (OR). P values for the strength of evidence for independent variables were assessed using likelihood ratio tests (LRT). Variables were assessed for inclusion in the multivariable model based on a criterion of *p*>0.2, conservatively set to ensure all potentially associated variables were included. Spearman’s rank tests were conducted on variables to observe correlation. To reduce high degrees of multicollinearity between independent variables, variance inflation factors (VIF) were examined following a stepwise procedure until only predictor variables meeting a moderate threshold of VIF≤6 remained in the global model (Rogerson, 2001). Variables were assessed with a backwards selection strategy until a predetermined threshold of α<0.05.

### Bioinformatics

Raw reads were assembled using SPAdes v 3.15.5 (Prjibelski *et al*., 2020) default parameters. The most likely contig corresponding to the mitochondrial genome was identified by searching the assembled contigs with BLASTn (Altschul *et al*., 1990), using the mitochondrial genome of *Trichuris muris* [NC_028621.1] as a query. Similarity of the extracted putative mitochondrial genome to published sequence data was established through BLASTn searches of all records including ‘Nematoda (taxid:6231)’. The putative mitochondrial genome was used as input for the MITOS annotation webserver (Bernt *et al*., 2013).

## Results

### Small mammal trapping success and diversity at different sites

Four villages were included in this study, selected from Mayuge District and Bugiri District in Eastern Uganda. Cross-sectional sampling (trapping over a single dry season) was conducted in 2 villages (Bugoto and Walumbe) in 2018, whilst in Musubi and Waka Waka repeat sampling was conducted over two consecutive dry seasons in 2018 and 2019. Traps were set in 98 participating households. 224 small mammals were trapped across the four villages over two dry seasons. Adjusted trap success by village and season ranged from 10.8% to 37.9% (Table 2). Small mammals were caught in 76.5% of households overall (N=75/98), ranging from 53.8% of households in Bugoto to 80.8% of households in Waka Waka in 2019. Musubi saw an increase in proportion of households with small mammals from 55.6% in 2018 to 77.3% in 2019. Each household trapped a mean of 1.6 small mammals, ranging from 0 to a maximum of 6 in a single house.

**Table 2.** Trapping data and adjusted trap success by site and session.

|  | Bugoto | Musubi |  | Walumbe | Waka Waka |  |
| --- | --- | --- | --- | --- | --- | --- |
|  | 2018 | 2018 | 2019 | 2018 | 2018 | 2019 |
| Sampling nights per village (n) | 5 | 5 | 3 | 3 | 3 | 3 |
| Total household trap nights per village (n) | 27 | 84 | 66 | 66 | 67 | 72 |
| Adjusted trap nights per village (n) | 29.0 | 347.0 | 276.5 | 284.5 | 289.0 | 292.0 |
| Total number full traps (n) | 11 | 38 | 30 | 46 | 35 | 51 |
| Adjusted trap success (%) | 37.9% | 11.0% | 10.8% | 16.2% | 12.1% | 17.5% |
| Total trapped (n) | 18 | 44 | 30 | 46 | 35 | 51 |
| Households with successful trapping (%) | 53.8% | 55.6% | 77.3% | 79.2% | 75.0% | 80.8% |
| Shannon Index (H) | 0.0 | 1.16 | 1.55 | 2.15 | 1.01 | 1.30 |
| Number of participating households | 13 | 27 | 22 | 24 | 24 | 26 |

Household level surveys were conducted to assess perceptions of household infestation levels (N=81). Nearly every household saw rodents in households, with many (71.6%, N= 58/71) reporting seeing rodents daily. Overall, 80.2% of households surveyed engaged in pest control (N=65/81), and 75.3% of households used chemical methods (N=61/81), consistent across villages (Table S2). Of households using chemical pest control, the most reported type was indometacin tablets, brand name Indocid (N=44/61). Other chemical methods include Fuko-Kil (N=3/61) and Rat-Rat (N=2/61) and unidentified rodenticide pastes or powders (N=15/61).

Postmortems and pathogen screening was conducted on 234 small mammals in total, including small mammals that were brought to the study in non-standard traps and cases of multiple trap occupancy (N=10) (Table 3). Of these, 50.4% were female (N=118/234) and 44.4% were male (N=104/234), with 12 individuals not sexed. The majority (73.1%; N=171/234) were classified as sexually mature based on external features. 135 of 234 were captured in Sherman traps (57.7%) compared to 87 in Tomahawk or Tomahawk-sized treadle traps (37.2%).

**Table 3.**
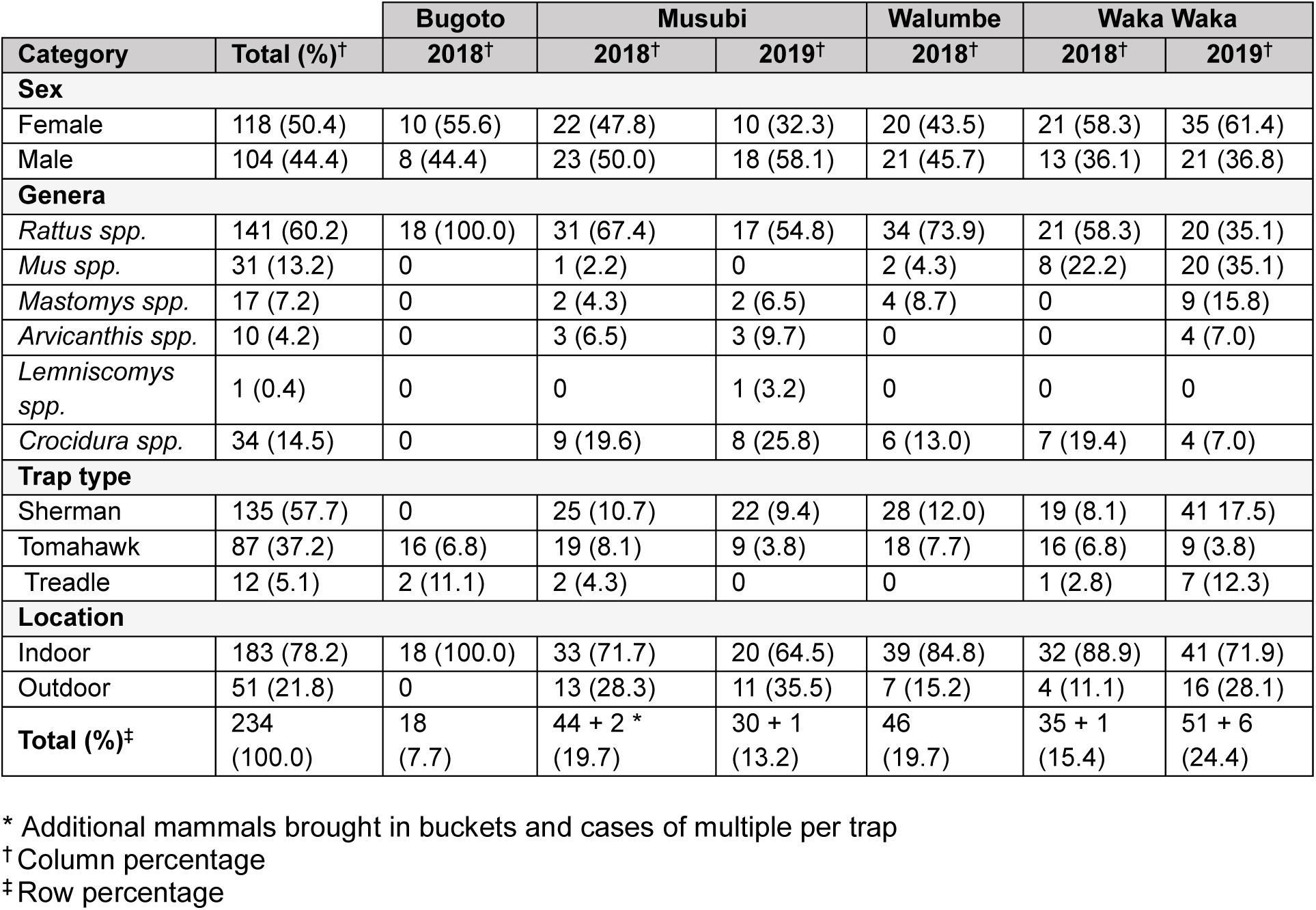
Summary of characteristics of rodents and shrews table **(N=234)**

|  |  | Bugoto | Musubi |  | Walumbe | Waka Waka |  |
| --- | --- | --- | --- | --- | --- | --- | --- |
| Category | Total (%)<sup>†</sup> | 2018<sup>†</sup> | 2018<sup>†</sup> | 2019<sup>†</sup> | 2018<sup>†</sup> | 2018<sup>†</sup> | 2019<sup>†</sup> |
| Sex |  |  |  |  |  |  |  |
| Female | 118 (50.4) | 10 (55.6) | 22 (47.8) | 10 (32.3) | 20 (43.5) | 21 (58.3) | 35 (61.4) |
| Male | 104 (44.4) | 8 (44.4) | 23 (50.0) | 18 (58.1) | 21 (45.7) | 13 (36.1) | 21 (36.8) |
| Genera |  |  |  |  |  |  |  |
| *Rattus spp.* | 141 (60.2) | 18 (100.0) | 31 (67.4) | 17 (54.8) | 34 (73.9) | 21 (58.3) | 20 (35.1) |
| *Mus spp.* | 31 (13.2) | 0 | 1 (2.2) | 0 | 2 (4.3) | 8 (22.2) | 20 (35.1) |
| *Mastomys spp.* | 17 (7.2) | 0 | 2 (4.3) | 2 (6.5) | 4 (8.7) | 0 | 9 (15.8) |
| *Arvicanthis spp.* | 10 (4.2) | 0 | 3 (6.5) | 3 (9.7) | 0 | 0 | 4 (7.0) |
| *Lemniscomys spp.* | 1 (0.4) | 0 | 0 | 1 (3.2) | 0 | 0 | 0 |
| *Crocidura spp.* | 34 (14.5) | 0 | 9 (19.6) | 8 (25.8) | 6 (13.0) | 7 (19.4) | 4 (7.0) |
| Trap type |  |  |  |  |  |  |  |
| Sherman | 135 (57.7) | 0 | 25 (10.7) | 22 (9.4) | 28 (12.0) | 19 (8.1) | 41 (17.5) |
| Tomahawk | 87 (37.2) | 16 (6.8) | 19 (8.1) | 9 (3.8) | 18 (7.7) | 16 (6.8) | 9 (3.8) |
| Treadle | 12 (5.1) | 2 (11.1) | 2 (4.3) | 0 | 0 | 1 (2.8) | 7 (12.3) |
| Location |  |  |  |  |  |  |  |
| Indoor | 183 (78.2) | 18 (100.0) | 33 (71.7) | 20 (64.5) | 39 (84.8) | 32 (88.9) | 41 (71.9) |
| Outdoor | 51 (21.8) | 0 | 13 (28.3) | 11 (35.5) | 7 (15.2) | 4 (11.1) | 16 (28.1) |
| Total (%)<sup>‡</sup> | 234 (100.0) | 18 (7.7) | 44 + 2 * (19.7) | 30 + 1 (13.2) | 46 (19.7) | 35 + 1 (15.4) | 51 + 6 (24.4) |

\* Additional mammals brought in buckets and cases of multiple per trap
<sup>†</sup> Column percentage
<sup>‡</sup> Row percentage

Captured small mammals belonged to six distinct genera, primarily rodents (*Rattus spp., Mus spp., Mastomys spp., Arvicanthis spp., Lemniscomys spp.)* as well as shrews (*Crocidura spp.*) (Table 2). Distribution of genera according to study site and trapping session is illustrated in Figure 2 to include all individuals. *Rattus rattus* was the most commonly trapped (N=141/234, 60.3%), followed collectively by shrews (*Crocidura spp.:* N=34/234, 14.4%) and house and pygmy mice (*Mus spp.:* N=31/123, 13.2%). Less common were multimammate mice (*Mastomys spp.:* 17/234, 7.3%) and unstriped grass mice (*Arvicanthis spp*.: N=10/234, 4.3%), and only one striped grass mouse (*Lemniscomys spp.:* N=1/234, 0.4%) was captured in this study. Because morphological identification of species is not possible for all genera in this region (particularly *Mastomys*, *Mus* and *Crocidura*; Monadjem et al 2015), we also amplified and sequenced mitochondrial genes to attempt species identification. There were incomplete or non-conclusive sequences for 14 individuals, the remainder were identified to species via morphological or molecular methods (Dataset 1).

**Figure 2.**
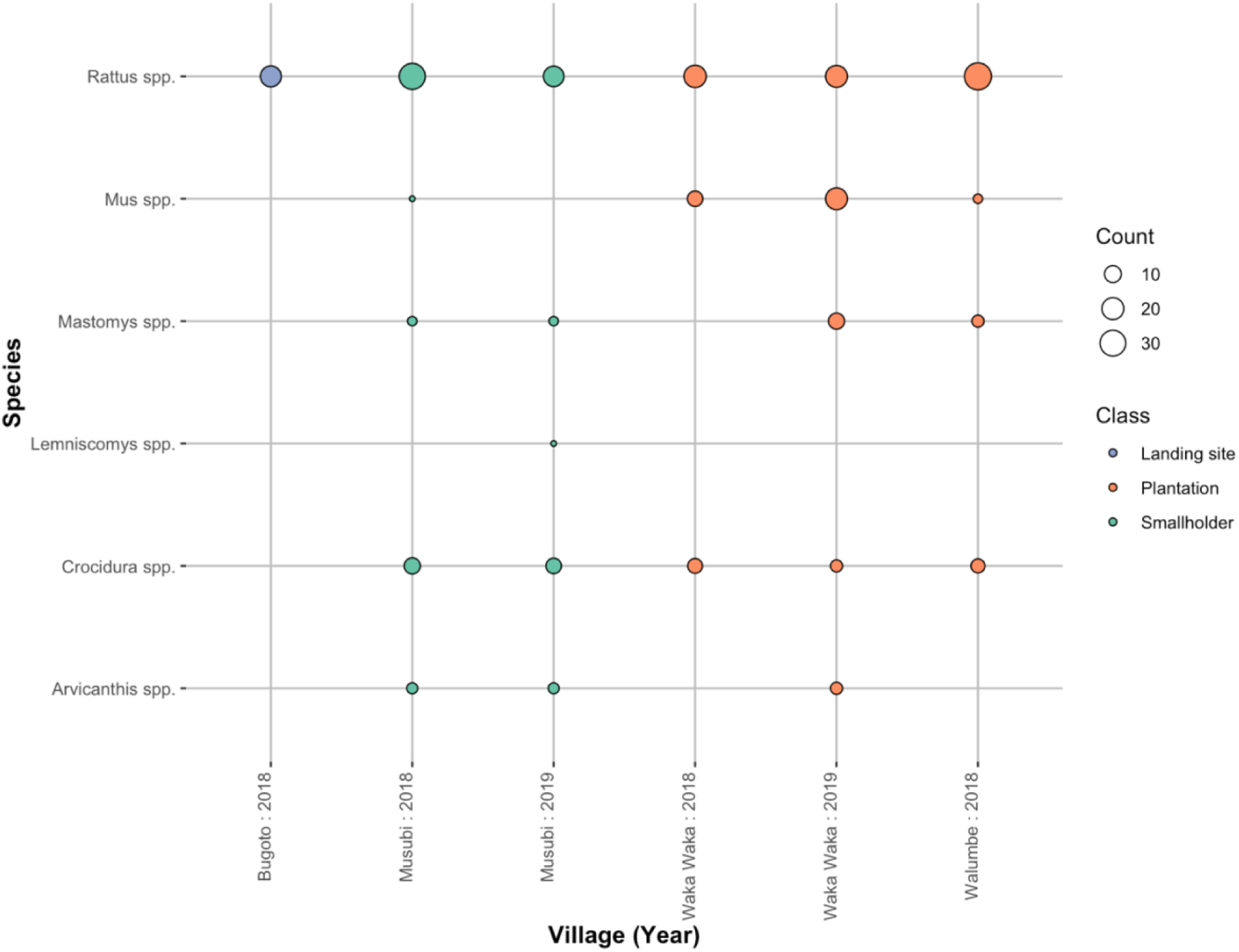
Bubble plot of small mammal genus and diversity across study site and year, coloured by land cover classification. **(N=234)**

Species accumulation curves were examined for overall sampling and disaggregated by village, with the caveat that some individuals were speciated and some were only identified to genus level. The plateau in the rarefication curve occurred at approximately 80 households and 15 species, indicating that increased intensity and stratification of sampling in each village and trapping season would be required to saturate species diversity (Supplementary Information, SI).

### *Village-level prevalence and distribution of* C. hepatica

Liver infection with *C. hepatica* was diagnosed both by gross parasitological examination and by PCR of both undiluted and a 1:10 dilution of liver DNA. Overall prevalence differed according to detection method (Table 4). Liver pathology was detected in 29% of small mammal postmortems (68/234, CI95% 23.2–34.9). PCR on undiluted DNA yielded an overall prevalence of 44.8% in a subset of small mammal samples that were tested using molecular methods (91/203, CI95% 38.0–51.7), compared to a lower overall prevalence of 26.1% (53/203, CI95% 20.0–32.2) using a 1:10 dilution. These prevalence estimates include faint bands, which are likely false positives. We considered a true positive to be a strong band on either PCR or a faint band on both PCRs. This overall prevalence was 34.5% (70/203, CI95% 27.9–41.0) (Table 4).

**Table 4.** Characteristics of rodents tested and number/percentage of confirmed C. hepatica infections. LP = liver pathology, X1 = PCR positive, X10 = PCR positive dilution, P = combined overall positive (1 strong or both faint)

|  |  | N | LP+ (%)<sup>†</sup> | N<sup>M</sup> <sup>‡</sup> | X1+ (%)<sup>†</sup> | X10+ (%)<sup>†</sup> | P+ (%)<sup>†</sup> | CI95%<sup>§</sup> |
| --- | --- | --- | --- | --- | --- | --- | --- | --- |
| **Genus** | *Arvicanthis spp.* | 10 | 6 (60.0) | 9 | 3 (33.3) | 5 (55.6) | 4 (44.4) | (12.0 – 76.9) |
|  | *Crocidura spp.* | 31 | 1 (2.9) | 29 | 12 (41.4) | 10 (34.5) | 8 (27.6) | (11.3 – 43.9) |
|  | *Lemniscomys spp.* | 1 | 0 (0.0) | 1 | 0 (0.0) | 0 (0.0) | 0 (0.0) | – |
|  | *Mastomys spp.* | 17 | 7 (41.2) | 15 | 7 (46.7) | 6 (40.0) | 9 (60.0) | (35.2 – 84.8) |
|  | *Mus spp.* | 20 | 4 (13.3) | 29 | 9 (31.0) | 2 (6.9) | 7 (24.1) | (8.6 – 39.7) |
|  | *Rattus spp.* | 142 | 50 (35.2) | 120 | 60 (50.0) | 30 (25.0) | 42 (35.0) | (26.5 – 43.5) |
| **Trap** | Sherman | 135 | 21 (15.5) | 124 | 50 (40.3) | 28 (22.6) | 38 (30.6) | (22.5 – 38.8) |
|  | Tomahawk * | 87 | 45 (51.7) | 68 | 38 (55.9) | 21 (30.9) | 30 (44.1) | (32.3 – 55.9) |
|  | Misc | 12 | 2 (16.7) | 11 | 3 (27.3) | 4 (36.4) | 2 (18.2) | (0.0 – 41.0) |
| **Location** | Indoor | 183 | 56 (30.6) | 159 | 72 (45.3) | 38 (28.9) | 50 (31.4) | (23.3 – 38.7) |
|  | Outdoor | 51 | 12 (23.5) | 44 | 19 (43.2) | 15 (34.1) | 20 (45.5) | (30.7 – 60.2) |
| **Site type** | Landing site | 18 | 3 (16.6) | 8 | 6 (75.0) | 4 (50.0) | 6 (75.0) | (45.0 – 100) |
|  | Plantation | 139 | 46 (33.1) | 127 | 51 (40.2) | 22 (17.3) | 34 (26.8) | (19.1 – 34.5) |
|  | Smallholder | 77 | 19 (24.7) | 68 | 34 (50.0) | 27 (39.7) | 30 (44.1) | (32.3 – 55.9) |
| **Village, Y** | Bugoto 2018 | 18 | 3 (16.7) | 8 | 6 (75.0) | 4 (50.0) | 6 (75.0) | (45.0 – 100) |
|  | Musubi 2018 | 46 | 15 (32.6) | 37 | 23 (62.2) | 21 (56.8) | 23 (62.2) | (46.5 – 77.8) |
|  | Musubi 2019 | 31 | 4 (12.9) | 31 | 11 (35.5) | 6 (19.4) | 7 (22.6) | (7.9 – 37.3) |
|  | Walumbe 2018 | 46 | 16 (34.8) | 41 | 19 (46.3) | 10 (24.4) | 11 (26.8) | (13.3 – 40.4) |
|  | Waka Waka 2018 | 36 | 13 (36.1) | 31 | 19 (61.3) | 6 (19.4) | 13 (41.9) | (24.6 – 56.3) |
|  | Waka Waka 2019 | 57 | 17 (29.8) | 55 | 13 (23.6) | 6 (10.9) | 10 (18.2) | (8.0 – 28.4) |
| **Overall** |  | 234 | 68 (29.1) | 203 | 91 (44.8) | 53 (26.1) | 70 (34.5) | (27.9 – 41.0) |

\* Tomahawk combined with Tomahawk-sized treadle traps
<sup>†</sup> Proportion of N positive for *C. hepatica* (row %)
<sup>‡</sup> Denominator (N<sup>M</sup>) small mammals tested by molecular methods
<sup>§</sup> 95% confidence interval (CI95%) calculated in R using count and sample size (binomial distribution)

Highest prevalence (overall combined positives) of *C. hepatica* in small mammals was observed in Bugoto, 2018, with 75% of small mammals tested by molecular methods found to be infected (6/8, CI95% 45.0–100.0). Lower prevalence was observed in Waka Waka in 2019, with 10/55 small mammals testing positive (18.2%, CI95% 8.0–28.4) and an intermediate proportion of small mammals infected in Walumbe, 2018 (26.8%, 11/41, CI95% 13.3–40.4). Prevalence was notably higher in sites categorised as landing sites at 75% (6/18, CI95% 45.0–100.0) compared to forest plantations areas (26.8%, 34/139, CI95% 19.9–34.5). Infection prevalence in small mammals in Musubi fell between consecutive trapping seasons, with high infection prevalence of 62.2% in 2018 (23/37, CI95% 46.5–77.8) and only 22.6% in the following trapping season (23/37, CI95% 7.9–37.3). In Waka Waka, a similar temporal trend was observed, though not found to be significant at a 95% confidence threshold (Figure 3). Prevalence of *C. hepatica* in small mammals does not appear to differ significantly between trapping location (inside/outside household), trap type or small mammal Genus. Differences between villages and trapping seasons is not reflected in categorical description of site, suggesting other landscape or ecological features is driving differences in infection burden.

**Figure 3.**
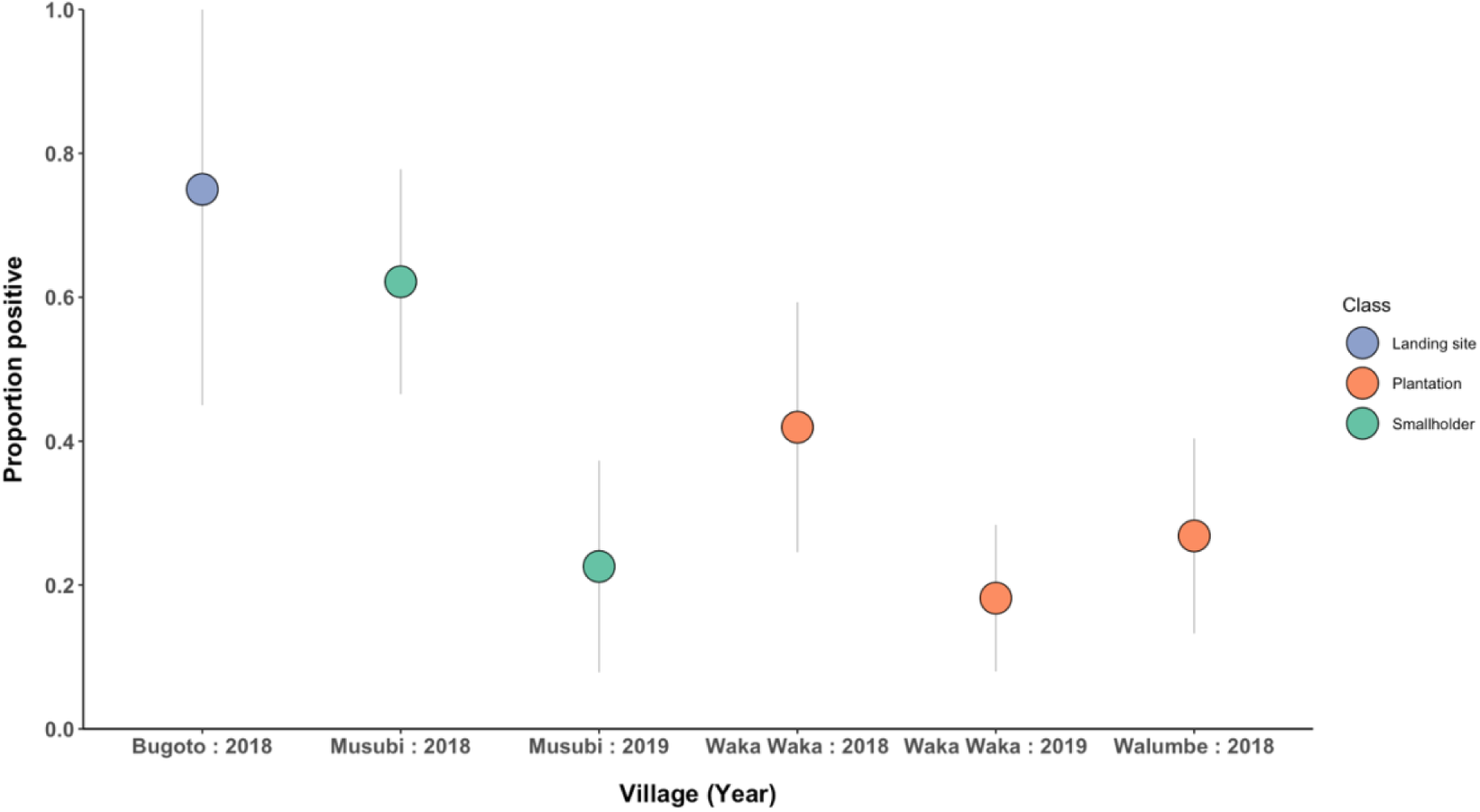
Proportion of small mammals infected by *C. hepatica* by village site and trapping season. Infection status was determined by molecular detection in one or two assays. Colour coded by site landscape classification.

### Landscape effects on village-level infection

Landscape effects were first evaluated with generalized linear mixed model regression to assess the association between individual village covariates and prevalence. A negative association between village level *C. hepatica* infection prevalence and mean monthly climatic factors (maximum and minimum temperature), as well as weak associations with human population density estimates and absolute forest cover was found (Figure S5). A strong negative association was observed between the ratio of other genera to *Rattus rattus*, a proxy for small mammal community diversity between villages and seasons, and *C. hepatica* prevalence. Following model selection, the best multivariable generalized linear mixed model regression included the ratio of other small mammal genera to Rattus rattus, building density, forest cover, and proportion of houses with rodents (Table 5). A significant strong negative association was retained between the ratio of all other small mammal genera to *Rattus rattus* and prevalence of *C. hepatica*.

**Table 5.** Multivariable generalized linear mixed model results for infection with *C. hepatica*.

|  | aOR | CI 95% |  | P value † |
| --- | --- | --- | --- | --- |
| Ratio (Other: *Rattus rattus*) | 0.45 | 0.26 | 0.77 | 0.003775 ** |
| Building density (count) | 0.86 | 0.221435 | 3.671188 | 0.7895 |
| Forest cover (count) * | 1.75 | 0.434412 | 10.40573 | 0.3906 |
| Proportion of houses with rodents (%) | 2.13 | 0.795394 | 5.41109 | 0.1129 |

Signif. codes: 0 '\*\*\*' 0.001 '\*\*' 0.01 '\*' 0.05 '.' 0.1 ' ' 1
\* Threshold for forest = 30% tree canopy cover
† Derived from likelihood ratio test (LRT)

### Mitogenome recovery

Sequencing of an adult worm recovered an assembly of mitochondrial DNA. BLASTn query of the *Trichuris muris* mitochondrial genome against the final assembly reported a single hit (NC_028621.1), of 4159bp in length. A BLASTn search of this hit against Genbank: Nucleotide records for Nematoda returned 58 hits (**Table S4**). The top four hits had query covers of 88%, 96%, 63% and 50%, e-values of 0, and corresponded to corresponded to *Pseudocapillaria tomentosa, Aonchotheca putorii, Capillaria sp cat-2108,* and *Eucoleus* annulatus, respectively, all parasitic nematodes in the *Capillaridae* family. The top (5th) non Capillariaid hit was to *Trichuris arvicolae* (query cover: 19%, e-value 0). MITOS annotation identified 36 gene, tRNA and rRNA features. The putative mitochondrial genome annotations with those of *Trichuris muris* [**acc**] and *Pseudocapillaria tomentosa* [**acc**]. The putative mitochondrial *C. hepatica* was identical in feature order to that of *P. tomentosa.* Missing features [**in b12_12**] with respect to *T. muris* and *P. tomentosa* were the *Atp8* gene, and tRNA-Tyr, as well as an interrupted 16sRNA annotation.

## Discussion

Land use change is known to disrupt ecological and social systems in ways that can affect pathogen dynamics. Parasite infection prevalence and intensity can be influenced by host community composition, host density, variation in susceptibility between host species or environmental features. We observe a heavy parasite burden of *C. hepatica*, with high prevalence across diverse species, both inside and outside residences, and evidence of village-level spatial heterogeneity in prevalence. We identified strong links between higher ratio of *Rattus rattus* to other native small mammal species to higher village-level prevalence. We also observed lower prevalence in small-holder agriculture. Preliminary results offer evidence of a dilution effect, with low-moderate canopy cover sufficient to support higher overall species diversity adjacent to small-holder agriculture and timber plantations. To our knowledge, this is the first study to screen synanthropic small mammals for *C. hepatica* using molecular methods in East Africa. These findings provide insight into ecological mechanisms of maintenance and transmission for a zoonotic pathogen in the context of changing land use and land cover and indicate the need for further molecular and ecological studies characterising *C. hepatica* and associated pathogens across a landscape gradient.

Infection prevalence and host abundance are key prerequisites for spillover into human populations. Despite consistently high coverage of chemical pest control, high levels of household infestation and incursion were recorded in all villages. We observed higher levels of household incursion in 2019 compared to 2018, but this was only in two sites. The temporal trend was consistent with loss in forest cover and the observation that proportion of forest cover is negatively correlated to household infestation (higher household infestation in areas with less forest cover). Although below thresholds for statistical significance, results are also indicative of an association between high household infestation (a metric for small mammal abundance) and higher prevalence, suggesting in this context that abundance may modulate prevalence in determining overall force of infection and have consequences for human risk. A heavily infected *Mastomys natalensis* was found in a home of a patient with hepatic calodiasis (Cochrane et al 1957). While human risk for infection is expected to be tightly linked with infestation and prevalence, more studies are needed to demonstrate if household infestation with infected small mammals is sufficient, or if quantifying transmission at larger spatial scales is required to inform human risk.

Across four sites and two dry seasons, we captured six genera: five rodent (*Mastomys, Rattus, Mus, Arvicanthis, Lemniscomys)* and one shrew (*Crocidura*). All sites recorded *Arvicanthis spp., Crocidura spp., Mastomys spp., Mus spp.* and *Rattus spp.,* except for Bugoto, where exclusively *Rattus rattus* was identified. The focus on households, and land immediately surrounding households may have limited the species we were able to detect. In the region there is a higher diversity of rodents in agricultural fields, including *Lophuromys* and *Dasymys* (Mayamba et al 2019), that were not included in this study due to sampling design. Adjusted trap success (ATS) showed a strong positive correlation to building density (ρ=0.83), indicating higher trap success in areas with dense infrastructure, and a strong negative correlation coefficient with Shannon Index (H) (ρ=-0.88) indicating higher trap success in sites dominated by fewer species. Bugoto is characterised as a ‘landing site,’ which denotes a village related to fishing trade, often urbanised, and densely populated. Bugoto exceeds the upper standard deviation for building density, and recent longitudinal studies suggest that a dense urban patch is likely to favour increased dominance of highly adapted species like *Rattus spp.* and local extirpation of other species via interspecific competition (Teitelbaum, 2024). It was also noted by inhabitants that when ‘stinky rats’ (a colloquialism for shrews, *Crocidura spp.*) are present, there is a noticeable absence of other rodent species. Whilst not a pattern that we observed in our data, this is supported by recent diet analysis and DNA metabarcoding of *Crocidura olivieri* that confirmed predation on *Mus musculus* (Galan *et al*., 2023).

Consistent with *C. hepatica* being a generalist nematode parasite, *C. hepatica* was observed in almost all species examined. Moderate infection prevalence was observed in native species: including *Mastomys spp.* (60.0%, CI95% 35.2–84.8) and African Giant White-toothed shrews (*Crocidura olivieri*, 27.6%, CI95% 11.3–43.9). Both species were found inside and outside households, suggesting they pose a risk to humans within households but can also link to small mammal assemblages in neighbouring fields and habitats. *C. hepatica* has previously been identified in shrews in Europe, USA and Asia (Li *et al*., 2010; Fuehrer, 2013; Miterpáková *et al*., 2024), but to our knowledge, this is the first evidence of infection in African shrews. The only host without *C. hepatica* was *Lemniscomys striatus*, but only one individual was checked in our survey. Overall, we found ubiquitous infection of *C. hepatica*, across all host genera and locations. The nematode parasite is common in this region in rodents and shrews, many of which are trapped within households, posing a risk for humans in these communities.

To identify infection with *C. hepatica*, we first used gross parasitological examination in the field and followed up with molecular diagnostics. Molecular detection, as expected, was higher than gross parasitological evaluation, but came with additional challenges. While the majority of individuals were screened molecularly, 13.2% of samples were omitted due to DNA degradation or sample loss during processing. We also had difficulties using published primers to consistently amplify *Calodium* DNA, which another study from Hong Kong (Manor et al. 2021) also found. In order to address this, we generated a mitogenome from a worm isolated in the study region using shot-gun sequencing and de novo assembly. The mitogenome is highly divergent from published mitochondrial genomes and can be used in the future to design better primers. In recent years, additional genetic sequences have been made available (Buńkowska-Gawlik et al. 2017, Tamaru et al. 2025), which will improve the development of sensitive and specific diagnostics for this understudied nematode.

Clear differences in village-level *C. hepatica* prevalence were observed between different sites. Land use and land cover heterogeneity is known to be linked to parasite infection prevalence in wildlife (Roberts *et al*., 1992; Gillespie and Chapman, 2008). We hypothesised that expansion of agriculture might disrupt ecological equilibrium of rodent assemblages and result in higher pathogen burden. Instead, we found no statistical relationship between village-level prevalence in rodents and recent forest loss, although it is important to note that the study was not designed to explicitly test for this relationship. Recent global meta-analyses find that habitat loss may not be a significant global driver of disease prevalence (Mahon *et al*., 2024) and only weak effects of forest loss on parasite prevalence (Heckley and Becker, 2025). Consensus dictates that parasite prevalence response to forest loss will be complex, context dependent, and contingent on parasite taxa and transmission mode (Gottdenker *et al*., 2014). However, by site description, prevalence was found to be significantly lower in sites adjacent to forest plantations (Waka Waka, Walumbe) compared to landing sites (Bugoto). Timber plantations may be affecting small mammal populations by supporting predators and act as ecological buffer against higher infection prevalence. While tree cover and canopy height provide an approximation of land cover composition meaningful for small mammal ecology, future work would be enriched by consideration of land cover configuration (connectivity, fragmentation, patch statistics) that influence rodent community diversity and abundance (Morales-Díaz *et al*., 2019) and pathogen dynamics (Teitelbaum, 2024).

Biodiversity and species community composition can have differential effects pathogen prevalence in a host population. In multivariable analyses, the only significant predictor of *C. hepatica* prevalence to explain village-level differences was ratio of native small mammal species to *Rattus rattus:* with higher ratio of native species to *Rattus rattus* (more diverse species assemblage), *C. hepatica* prevalence is lower. Theoretical frameworks show that in some contexts, higher species diversity (higher Shannon Index, H) is associated with higher intensity transmission but decreases in overall proportion of infectious individuals (lower overall prevalence) (Roche *et al*., 2012). Empirical longitudinal data on land consolidation and rodent-pathogen dynamics provides a link between rodent ecology, disease ecology and land use, with evidence that in addition to favouring a single dominant species (lower H), land consolidation results in higher pathogen prevalence in the newly dominant host species (Teitelbaum, 2024, Pei et al 2024). Likewise, in Sierra Leone and Guinea, presence of *Rattus rattus* decreased prevalence of Lassa virus by competing with the dominant reservoir *Mastomys natalensis* (Eskew *et al*., 2024). In this system, dominance of generalist species *Rattus rattus* may potentiate *C. hepatica* burden through interspecies competition, while native species – supported by more heterogenous vegetation – creates a dilution effect (Keesing *et al*., 2006). However, this study only trapped in four villages with two time-points, limiting the statistical power to infer meaningful relationships between environmental covariates and village-level differences in rodent abundance or parasite prevalence. In future analyses, more robust estimates of population prevalence could have been obtained with hidden Markov models to account for a latent observation process. Individual variation in pathogen load was not captured by the scope of the study. Understanding how local land use changes species composition and competition will be key to predicting pathogen burden.

## Conclusion

Infection prevalence and host abundance play critical roles in the dispersal and transmission of disease. Prevalence and intensity of parasite infection in wildlife hosts are affected by the environment, host density, and individual characteristics. Using both microscopic and molecular diagnostics, we confirm a broad host range, with *C. hepatica* in all sites and in all invasive and native species of rodents and shrews. Building density appears to potentiate conditions for *C. hepatica,* with both high abundance of *Rattus rattus* and high parasite prevalence found in the densest human settlements, a concern for further exacerbation of zoonotic risk in vulnerable communities. Future work should focus on understanding key transmission routes for *C. hepatica* and specific hazard to public health.

## Data availability

Data on household trap success and details of postmortems are available as .csv files from Enlighten. Sequence data from host () and parasites (PX206072) is deposited on GenBank.

## Acknowledgements

We appreciate the many community members and leaders that allowed us to conduct surveys in their homes and villages. We also appreciate the Vector Control Division Research Ethics Committee that evaluated the protocol and came out to monitor activities and provide feedback. This work could not have happened without the support of Village Health Teams and the Vector Control Division’s strong public health network.

## Author’s contribution

CF and KA conceived and designed the study. TD and CF performed bioinformatics. KA trained field personnel. CLF, DA, MA, MA AN, RB, JR and OE performed fieldwork. MC, CLF, DA, MA, CR, RB, FB and OE performed laboratory work. CLF wrote the first draft with contributions from TD and MC. EJ conducted the statistical data analysis and compiled the final draft.

## Ethical standards

All trapping protocols and household surveys are covered by ethical approval from the Vector Control Research Ethics Committee (VCDREC-095), the Ugandan National Council for Science and Technology (NS-633) and the School of Veterinary Medicine at the University of Glasgow (Ref. 27a/18).

