## Supplementary Information for "Prevalence of zoonotic hepatic nematode varies with small mammal community diversity across a heterogenous landscape in Eastern Uganda"

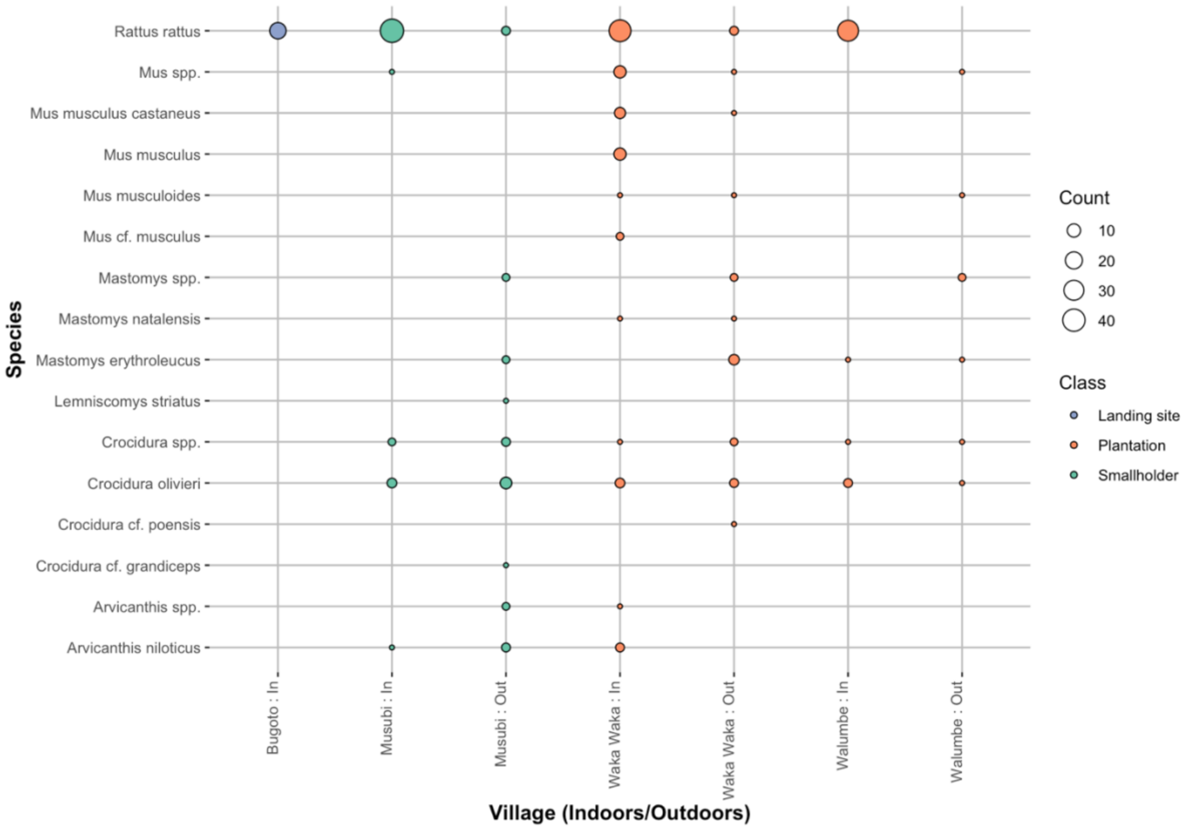


**SI Figure 1:** Bubble plot of species abundance by minimum classification of species/genera, village and whether small mammal was trapped inside or outside the household. Colour coded by land class.

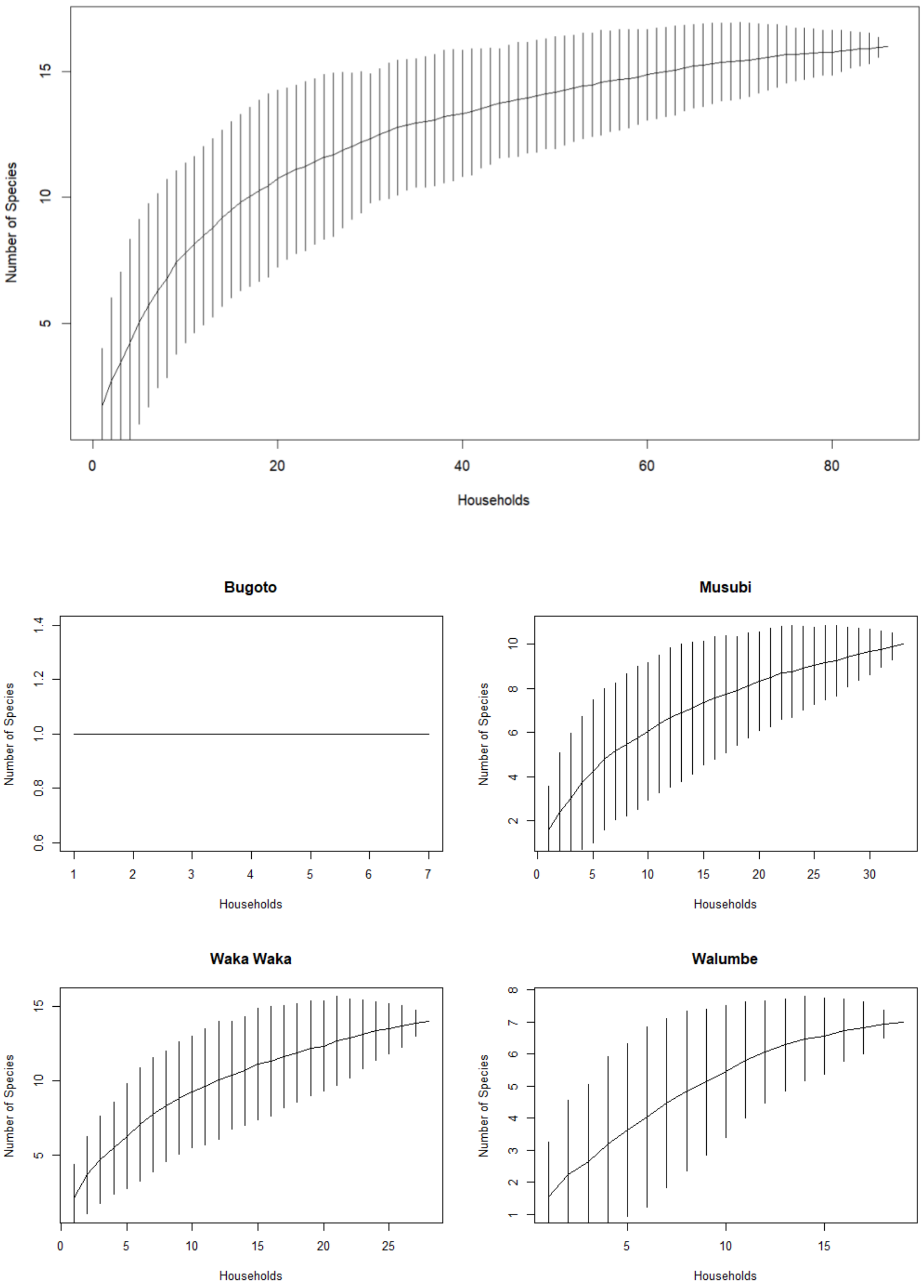


**SI Figure 2. A.** Species accumulation (rarefication curve) for all species (minimum disaggregation) and all households over two sampling periods. **B.** Rarefication curves per village site. In Bugoto only *Rattus rattus* was trapped, resulting in a flatline curve of 1.

**Table S1:** Mean and distribution (SD) of environmental characteristics per village centroid within 0.5km buffer radii

| | | Temp. scale | Bugoto | Musubi | Waka Waka | Walumbe | Mean | SD |
|---|---|---|---|---|---|---|---|---|
| Elevation (m) | | 2000 | 1140.32 | 1145.84 | 1141.15 | 1141.62 | 1142.23 | 2.47 |
| Temp. min (Feb) (°C) | | 1970-2000 | 16.50 | 16.50 | 16.50 | 16.30 | 16.45 | 0.10 |
| Temp. min (Aug) (°C) | | 1970-2000 | 15.90 | 15.90 | 15.80 | 15.80 | 15.85 | 0.56 |
| Temp. max (Feb) (°C) | | 1970-2000 | 29.60 | 29.60 | 29.70 | 29.30 | 29.55 | 0.18 |
| Temp. max (Aug) (°C) | | 1970-2000 | 27.20 | 27.20 | 27.11 | 27.10 | 27.15 | 0.06 |
| Population density (p/km$^2$) | | 2019 | 824.03 | 881.06 | 669.54 | 260.46 | 658.77 | 280.17 |
| Building density (count) | | 2023 | 730 | 213 | 502 | 513 | 489.50 | 212.13 |
| Percentage cropland (%) | | 2019 | 19.04 | 47.25 | 51.87 | 15.93 | 33.52 | 18.66 |
| Average forest height (m) | | 2019 | 28.23 | 11.86 | 24.91 | 35.26 | 25.06 | 9.80 |
| Forest cover (count) | 50% | 2009 | 108 | 129 | 22 | 136 | 98.75 | 52.53 |
| *Classification threshold* | 40% | 2009 | 209 | 233 | 38 | 191 | 188 | 107.92 |
| *% canopy cover per pixel* | 30% | 2009 | 344 | 323 | 63 | 235 | 241.25 | 127.87 |
| | 50% | 2014 | 108 | 129 | 18 | 136 | 97.75 | 54.48 |
| | 40% | 2014 | 290 | 233 | 32 | 191 | 186.50 | 110.70 |
| | 30% | 2014 | 344 | 323 | 57 | 235 | 239.75 | 130.66 |
| | 50% | 2017 | 108 | 129 | 18 | 136 | 97.75 | 54.48 |
| | 40% | 2017 | 290 | 233 | 32 | 191 | 186.50 | 110.70 |
| | 30% | 2017 | 344 | 323 | 57 | 235 | 239.75 | 130.66 |
| | 50% | 2019 | 108 | 129 | 18 | 136 | 97.75 | 54.48 |
| | 40% | 2019 | 290 | 230 | 32 | 191 | 185.75 | 110.29 |
| | 30% | 2019 | 344 | 318 | 57 | 235 | 238.5 | 129.52 |
| Forest cover (proportion) | 50% | 2009 | 0.1060 | 0.1265 | 0.0216 | 0.1337 | 0.10 | 0.05 |
| *Classification threshold* | 40% | 2009 | 0.2846 | 0.2284 | 0.0373 | 0.1878 | 0.18 | 0.11 |
| *% canopy cover per pixel* | 30% | 2009 | 0.3376 | 0.3167 | 0.0619 | 0.2311 | 0.24 | 0.13 |
| | 50% | 2014 | 0.1060 | 0.1265 | 0.0177 | 0.1337 | 0.10 | 0.05 |
| | 40% | 2014 | 0.2846 | 0.2284 | 0.0314 | 0.1878 | 0.18 | 0.11 |
| | 30% | 2014 | 0.3376 | 0.3167 | 0.0560 | 0.2311 | 0.24 | 0.13 |
| | 50% | 2017 | 0.1060 | 0.1265 | 0.0177 | 0.1337 | 0.10 | 0.05 |
| | 40% | 2017 | 0.2846 | 0.2284 | 0.0314 | 0.1878 | 0.18 | 0.11 |
| | 30% | 2017 | 0.3376 | 0.3167 | 0.0560 | 0.2311 | 0.24 | 0.13 |
| | 50% | 2019 | 0.1060 | 0.1265 | 0.0177 | 0.1337 | 0.10 | 0.05 |
| | 40% | 2019 | 0.2846 | 0.2255 | 0.0314 | 0.1878 | 0.18 | 0.11 |
| | 30% | 2019 | 0.3376 | 0.3118 | 0.0560 | 0.2311 | 0.23 | 0.13 |

**Table S2:** Summary of results from household rodent/small mammal survey (**N=81**)

| | | **N (%)** |
|---|---|---|
| **Rodents sighted** | Every day | 58 (71.6) |
| | More than once a week | 4 (4.9) |
| | Less than once a week | 16 (19.8) |
| | NA | 3 (3.7) |
| **Rodent evidence in house** | Every day | 57 (70.4) |
| | More than once a week | 7 (8.6) |
| | Less than once a week | 15 (18.5) |
| | NA | 2 (2.5) |
| **Rodent evidence in surroundings** | Every day | 40 (49.4) |
| | More than once a week | 9 (11.1) |
| | Less than once a week | 24 (29.6) |
| | Never | 6 (7.4) |
| | NA | 2 (2.5) |
| **Rodent evidence in garden** | Every day | 12 (14.8) |
| | More than once a week | 26 (32.1) |
| | Less than once a week | 21 (25.9) |
| | Never | 6 (7.4) |
| | No fields | 13 (16.0) |
| | NA | 3 (3.7) |
| **Rodent evidence in kitchen** | Every day | 49 (60.5) |
| | More than once a week | 10 (12.3) |
| | Less than once a week | 19 (23.5) |
| | Never | 1 (1.2) |
| | NA | 2 (2.5) |
| **Garden location** | Adjacent/less than 5 min walk | 30 (37.0) |
| | More than 5 min walk | 37 (45.7) |
| | NA | 14 (17.3) |
| **Pest control** | Yes | 65 (80.2) |
| | No | 14 (17.3) |
| | NA | 2 (2.5) |
| **Pest control type** | Chemical | 61 (75.3) |
| | Mechanical | 4 (4.9) |
| | NA | 16 (19.6 |
| **Total** | | **81 (100%)** |

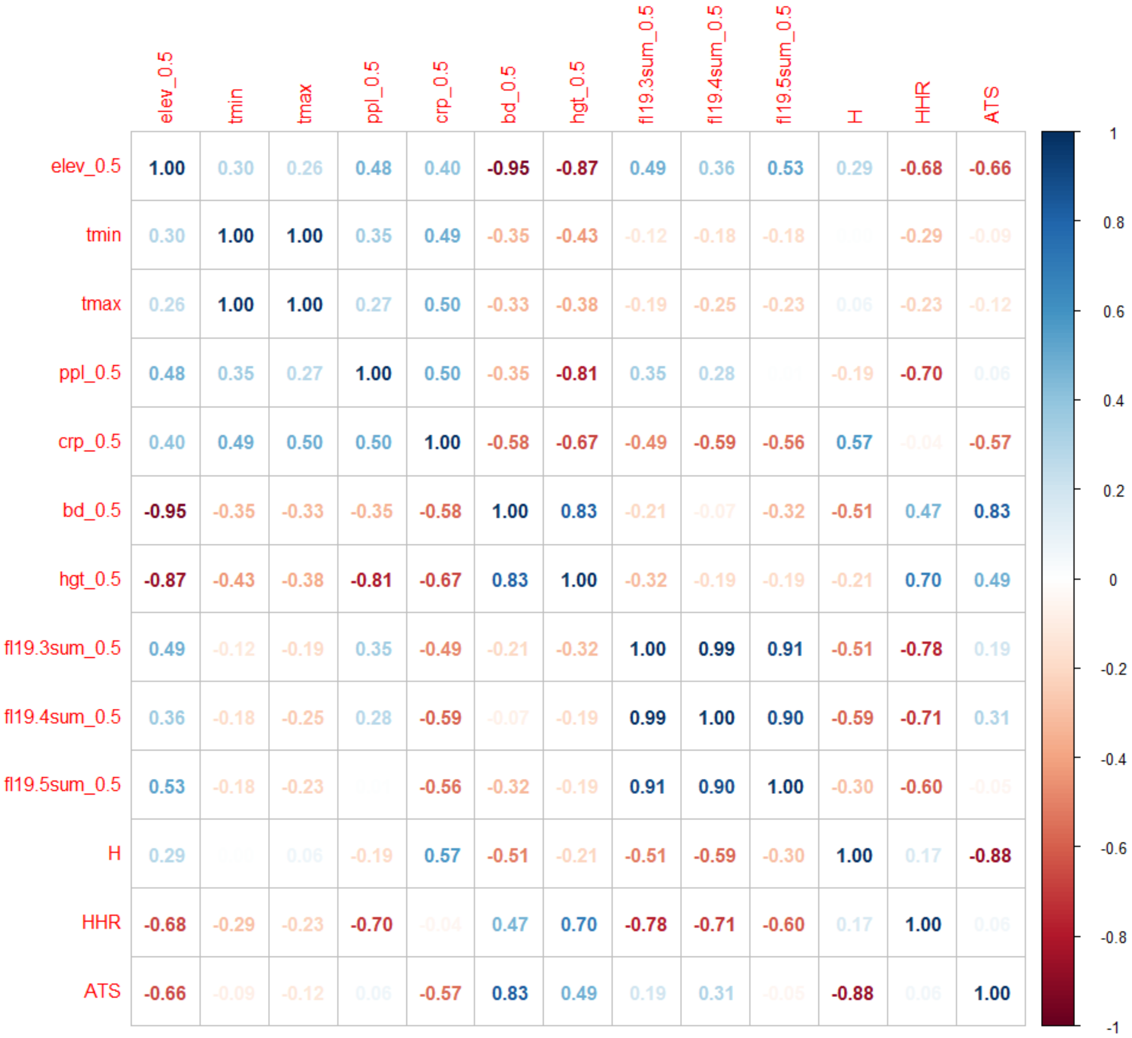


**Figure S3:** Spearman's rank correlation matrix of environmental and ecological variables remaining after bivariable selection. H = Shannon Index for diversity; HHR = household rattiness (%); ATS = adjusted trap success.

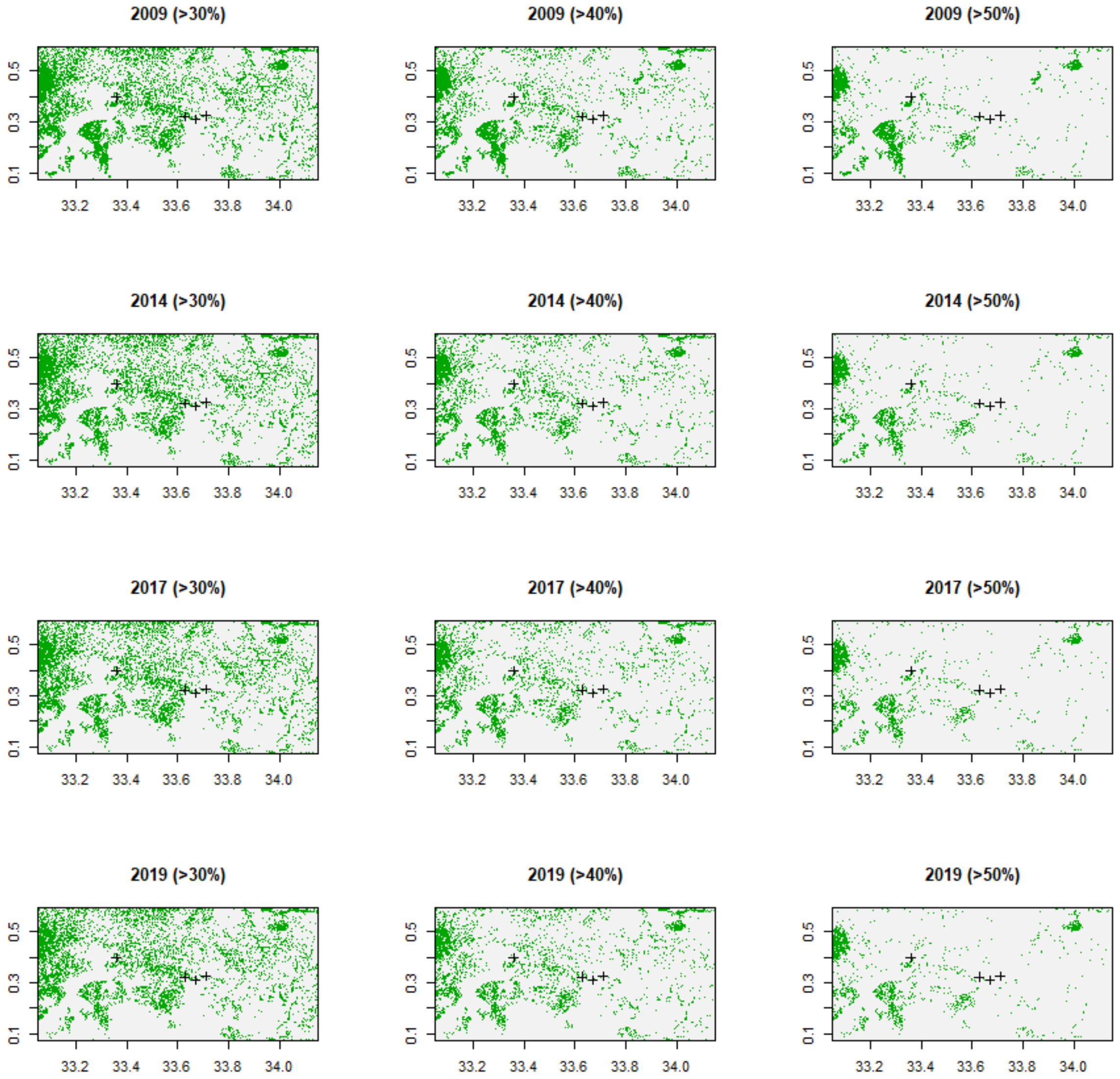


**Figure S4**: Forest (green) and non-forest (white) progressively lost in years 2009, 2014, 2017 and 2019 (plots top to bottom). Plots left to right show increasingly conservative definition of forest, according to proportion of canopy cover (>30%, >40% and >50% most conservatively).

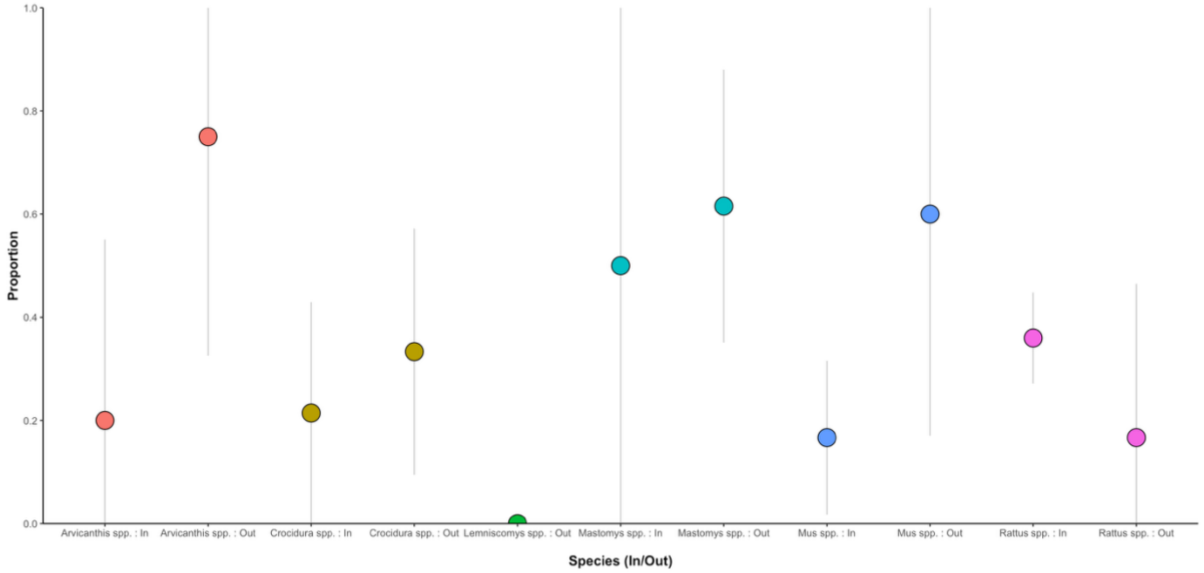


**Figure S5.** Proportion of small mammals infected with *C. hepatica* (according to faint signal on both diluted and undiluted DNA PCR or strong signal on one diagnostic) by species and trapping location inside or outside households. Colour coded by species.

**Table S3.** Bivariable binomial generalised linear regression analysis of effect of landscape and ecological covariates on village level prevalence of *C. hepatica.*

| | Coefficient | CI 95% | | P value † | |
|---|---|---|---|---|---|
| Elevation (m) | 1.127 | 0.99 | 1.28 | 0.0739 | . |
| Temperature min (°C) | 0.853 | 0.77 | 0.935 | 0.000911 | *** |
| Temperature max (°C) | 0.954 | 0.93 | 0.978 | 0.000311 | *** |
| Population density (p/km$^2$) | 1.00 | 1.00 | 1.00 | 0.03802 | * |
| Building density (count) | 1.00 | 1.00 | 1.00 | 0.245 | - |
| Percentage of cropland (%) | 0.78 | 0.11 | 5.95 | 0.807 | |
| Average forest height (m) | 0.97 | 0.94 | 1.00 | 0.068 | . |
| Forest cover (count) * | 1.00 | 1.00 | 1.01 | 0.0109 | * |
| Shannon Index (H) | 0.81 | 0.43 | 1.53 | 0.521 | |
| Proportion of houses with rodents (HHR) (%) | 0.99 | 0.97 | 1.00 | 0.123 | - |
| Ratio (Other: *Rattus rattus*) | 0.40 | 0.22 | 0.67 | 0.000961 | *** |
| Adjusted trap success (ATS) (%) | 0.99 | 0.94 | 1.03 | 0.567 | |

Signif. codes: 0 '***' 0.001 '**' 0.01 '*' 0.05 '.' 0.1 ' ' 1

* Threshold for forest = 30% tree canopy cover

† Derived from likelihood ratio test (LRT)

**Table S4. Top matches to mitochondrial DNA**

| scientificname | sseqid | pident | length | mismatch | evalue | bitscore |
|---|---|---|---|---|---|---|
| Pseudocapillaria tomentosa | MZ708958.1 | 78.091 | 9124 | 1631 | 0 | 5435 |
| Aonchotheca putorii | MZ708958.1 | 75.242 | 2892 | 596 | 0 | 1264 |
| Capillaria sp. cat-2018 | MZ708958.1 | 77.064 | 763 | 97 | 6.82E-98 | 368 |
| Eucoleus annulatus | NC_071371.1 | 79.085 | 5508 | 966 | 0 | 3618 |
| Trichuris arvicolae | NC_071371.1 | 81.342 | 3859 | 592 | 0 | 3020 |
| Trichuris muris | NC_071371.1 | 75.885 | 4603 | 862 | 0 | 2122 |
| Trichuris sp. 2 ARS-2017 | MH665363.1 | 73.934 | 6119 | 1094 | 0 | 2001 |
| Aonchotheca putorii | MH665363.1 | 75.508 | 3197 | 657 | 0 | 1450 |
| Trichuris sp. ETH232 | NC_056391.1 | 73.763 | 3617 | 774 | 0 | 1260 |
| Trichuris sp. KE396 | NC_056391.1 | 72.851 | 3757 | 752 | 0 | 1040 |
| Trichuris mastomysi | MZ229684.1 | 73.329 | 2857 | 595 | 0 | 900 |
| Trichuris suis | NC_028621.1 | 72.785 | 3013 | 634 | 0 | 850 |
| Trichuris muris | MZ229685.1 | 72.214 | 2836 | 602 | 0 | 701 |
| Calodium hepaticum | MZ229685.1 | 70.963 | 644 | 143 | 3.53E-21 | 113 |
| Trichuris sp. ETH392 | OP363931.1 | 84.356 | 652 | 102 | 1.31E-179 | 640 |
